# Sensorimotor peak alpha frequency is a reliable biomarker of pain sensitivity

**DOI:** 10.1101/613299

**Authors:** Andrew J. Furman, Mariya Prokhorenko, Michael L. Keaser, Jing Zhang, Shuo Chen, Ali Mazaheri, David A. Seminowicz

## Abstract

Previous research has observed that individuals with chronic pain demonstrate slower alpha band oscillations (8-12 Hz range) during resting electroencephalography (EEG) than do age-matched, healthy controls. While this slowing may reflect pathological changes within the brain that occur during the chronification of pain, an alternative explanation is that healthy individuals with slower alpha frequencies are more sensitive to prolonged pain, and by extension, more susceptible to developing chronic pain. To formally test this hypothesis, we examined the relationship between the pain-free, resting alpha frequency of healthy individuals and their subsequent sensitivity to two experimental models of prolonged pain, Phasic Heat Pain and Capsaicin Heat Pain, at two testing visits separated by 8 weeks on average (n = 61 Visit 1, n = 46 Visit 2). We observed that the speed of an individual’s pain-free alpha oscillations was negatively correlated with sensitivity to both prolonged pain tests and that this relationship was reliable across short (minutes) and long (weeks) timescales. Furthermore, we used the speed of pain-free alpha oscillations to successfully identify those individuals most sensitive to prolonged pain, which we also validated on data from a separate, independent study. These results suggest that alpha oscillation speed is a reliable biomarker of prolonged pain sensitivity with the potential to become a tool for prospectively identifying pain sensitivity in the clinic.

## Introduction

Chronic pain is a debilitating condition with cognitive, affective, and sensory symptoms that afflicts nearly one fifth of the American population (Kennedy et al., 2014), leading to treatment and work loss costs totaling nearly six hundred billion dollars annually (Gaskin & Richard, 2012). Identifying individuals at high risk for developing chronic pain is a crucial, but under-explored, avenue for combatting chronic pain and its related economic burdens. At present, prediction of chronic pain development is poor: for example, one of the best predictors of persistent post-surgical pain is the intensity of pain reported directly after surgery (e.g Katz et al., 1996). While useful for post-operative case management, these measures cannot be used to identify, and target prophylactic treatments to, individuals at risk for developing chronic pain. What is urgently needed is a measure of an individual’s sensitivity to prolonged pain that can be obtained prior to medical intervention. To that end, the objective of the current study is to systematically investigate the hypothesis that an individual’s peak alpha frequency, measured with resting state electroencephalography (EEG), is a trait-like marker of their sensitivity to prolonged pain.

The alpha rhythm (8-12 Hz) is the predominant oscillatory activity observed in scalp-recorded EEG of the primary sensory cortices (e.g. occipital, somatosensory) while an individual is quietly resting. Across individuals, there is considerable variability in the alpha band frequency from which the greatest power is recorded (Haegens et al, 2014, Bazanova & Vernon, 2014). This frequency, often labeled the Peak or Individual Alpha Frequency (PAF/IAF), has been suggested to contribute to individual differences in multiple psychological and physiological processes (e.g. Klimesch, 2012; Samaha and Postle, 2015; Gulbnaite et al., 2017; Mierau et al., 2017; Van Diepen et al., 2019).

Previous research has consistently observed abnormally slow PAF in chronic pain patients (Sarnthein et al., 2005; Walton et al., 2010; Lim et al., 2016), with increasingly slower PAF associated with increasingly longer durations of chronic pain (de Vries et al., 2013). This apparent slowing of PAF in chronic pain has been interpreted to reflect pathological changes within the brain that occur during the chronification of pain (Llinás et al., 1999). Work from our lab has, however, shown that slow PAF, recorded in the absence of pain (i.e. pain-free PAF), also reflects heightened sensitivity to prolonged pain in healthy individuals (Furman et al, 2018 & 2019). Given that heightened pain sensitivity is a risk factor for developing chronic pain (Diatchenko et al., 2005), an alternative interpretation of the aforementioned chronic pain findings is that slow PAF reflects an increased sensitivity to prolonged pain that predates disease onset. Put another way, slow PAF may reflect a predisposition for developing chronic pain rather than a result of its development.

In the current study, we sought to further characterize the relationship of pain-free PAF to prolonged pain sensitivity by exposing participants to two experimental models of prolonged pain, Phasic Heat Pain (PHP) and Capsaicin Heat Pain (CHP), at two testing visits separated by multiple weeks. This study design allowed us to test two key predictions of the hypothesis that pain-free PAF is a trait-like marker of an individual’s sensitivity to prolonged pain: (1) that pain-free PAF reflects pain sensitivity to multiple prolonged pain tests; and (2) that an individual’s pain-free PAF can predict their sensitivity to prolonged pain at more than one point in time. In addition to these main aims, and with an eye towards its potential clinical application, we also examined whether pain-free PAF can be used to successfully identify high and low pain sensitive individuals.

## Materials and methods

### Participants

Sixty-one pain-free, neurotypical adult participants (31 males, mean age = 27.82, age range = 21-42) took part in the experiment between 7/6/2016 and 10/20/2017. This study was approved by the University of Maryland, Baltimore Institutional Review Board, and informed written consent was obtained from each participant prior to any study procedures. The study was pre-registered on ClinicalTrials.gov (NCT02796625).

Table 1 provides information regarding how many participants contributed data to each analysis.

**Table 1.** Summary of exclusions and participants contributing data at each testing visit.

|  | Visit 1 |  |  | Visit 2 |  |  |
| --- | --- | --- | --- | --- | --- | --- |
|  | Capsaicin Responder | Capsaicin Non-Responder | High Pain Tolerance | Capsaicin Responder | Capsaicin Non-Responder | High Pain Tolerance |
| Total Participants | 35 (19 Female) | 15 (6 Female) | 11 (5 Female) | 27 (14 Female) | 11 (4 Female) | 8 (3 Female) |
| Exclusions |  |  |  |  |  |  |
| EEG Technical Error | 1 (Female) | 0 | 0 | 0 | 0 | 0 |
| Abnormal Pain Ratings | 1 (Female) | 0 | 1 (Male) | 1 (Female) | 0 | 1 (Male) |
| Abnormal CHP Change | 1 (Female) | 0 | 0 | 1 (Female) | 0 | 0 |
| Participants Remaining | 32 (16 Female) | 15 (6 Female) | 10 (5 Female) | 25 (12 Female) | 11 (4 Female) | 7 (3 Female) |

### EEG

Scalp EEG was collected from an EEG cap housing a 63 channel BrainVision actiCAP system (Brain Products GmbH, Munich, Germany) labeled according to an extended international 10–20 system (Oostenveld and Praamstra, 2001). All electrodes were referenced online to the average across all recording channels and a common ground set at the AFz site. Electrode impendences were maintained below 5 kΩ throughout the experiment. Brain activity was continuously recorded within a 0.01–100 Hz bandpass filter, and with a digital sampling rate of 500 Hz. The EEG signal was amplified and digitized using an actiCHamp DC amplifier (Brain Products GmbH, Munich, Germany) linked to BrainVision Recorder software (version 2.1, Brain Products GmbH, Munich, Germany).

### Thermal Stimulator and Pain Scale

Thermal stimuli were delivered to the volar surface of the participant’s left forearm using a thermal contact heat stimulator (27mm diameter Medoc Pathway CHEPS Peltier device; Medoc Advanced Medical Systems Ltd., Ramat Yishai, Israel).

Unless otherwise stated, pain ratings were collected continuously with a manual analog scale consisting of a physical sliding tab (Medoc Advanced Medical Systems Ltd., Ramat Yishai, Israel). Prior to testing, participants were instructed that the lower and upper bounds of the scale represented no pain and the most pain imaginable, respectively, and that they should continuously update the position slider to indicate the amount of pain currently being experienced. Care was taken by experimenters to avoid providing numerical anchors when describing the scale and no additional physical landmarks were present on the scale. Prior to testing, participants were given an opportunity to practice using the device with their eyes open and closed. During testing, participants were permitted to briefly open their eyes while rating.

Pain ratings were collected from the manual analog scale at a rate of 1000 Hz. Manual analog scale data was transformed by converting the horizontal position of the slider into a continuous value between 0 and 100.

### Quantitative Sensory Testing

Participants were asked to complete four threshold tests: 1) to report when they felt a temperature increase (Warmth Detection Threshold); (2) to report when they felt a temperature decrease (Cool Detection Threshold); (3) to report when an increasing temperature first became painful (Heat Pain Threshold); and (4) to report when a decreasing temperature first became painful (Cold Pain Threshold). A total of three trials were presented for each test with an ISI of 4-6 seconds (randomly determined on a per trial basis). Participants provided feedback for each test by clicking either the left or right button of a computer mouse placed in their right hand. For each test, temperatures were applied with a rise rate of 1°C/second and return rate of 2°C/second (initiated on any mouse click).

All testing was performed on the volar surface of the left forearm. The distance from the wrist to elbow joint was measured and the forearm was divided into three equal length zones. For each test, the first trial was administered to the zone closest to the wrist, the second trial administered to the middle forearm zone, and the third trial administered to the zone closest to the elbow.

### Phasic Heat Pain (PHP) Model

Temperatures used during the PHP model were determined during each participant’s initial screening visit to the laboratory (Visit 0). During these sessions, participants were exposed to 12, 20 second trials in which a single temperature (2.5 second rise and fall) was applied to the volar surface of the left forearm. At the conclusion of each trial, participants reported the average pain they experienced during temperature application; participants were instructed to report pain ratings on a scale of 0-10, with 0 indicating no pain and 10 the most pain imaginable. Temperatures ranged from 37 to 48°C (intervals of 2°C, starting as if 37°C was 38°C) and each temperature was presented twice in a pseudorandom order. Trials were separated by 10 seconds and after each trial the thermode was moved to a neighboring forearm zone in order to minimize sensitization. Using pain reports from these trials, the temperature that most closely evoked an average pain rating of 5/10 was selected. This level of pain was targeted in order to best match the intensity of pain evoked by the CHP model (Furman et al., 2018). For a few participants, none of temperatures were able to evoke a 5/10 pain rating. For these individuals, 48°C was used during PHP testing.

The PHP model itself consisted of a series of five consecutive stimulus trains. Each train lasted one minute and consisted of application of a predetermined temperature for 40 seconds (rise and fall times of 2s) followed by application of a neutral skin temperature stimulus (32°C) for 20 seconds. PHP scores were calculated by averaging pain ratings from the five, forty second periods in which the temperature was present.

### Capsaicin Heat Pain (CHP) Model

The CHP model lasts for hours to days and recapitulates some cardinal sensory aspects of chronic neuropathic pain (Culp et al., 1989; LaMotte RH, et al., 1992; Baron 2009; Lötsch et al., 2015) without causing lasting tissue damage (Henriques & Moritz, 1947). CHP procedures were similar to those used in our prior study (Furman et al., 2018). In brief, we applied ~1 g 10% capsaicin paste (Professional Arts Pharmacy, Baltimore, MD) topically to the volar surface of the left forearm, fixing it in place with a Tegaderm bandage. A thermode was then placed over top of the capsaicin application, heated to 40°C and held in place for 20 minutes to allow for capsaicin incubation. Given that pain from topically applied capsaicin varies as a function of skin temperature (Anderson et al., 2002), the thermode temperature was held at 40°C for all participants. This temperature was selected because, in the absence of capsaicin, most individuals find it non-painful thereby providing comfort that any pain generated by this temperature during capsaicin exposure is likely a consequence of the agent’s sensitizing effects. CHP scores were calculated by averaging ratings across the entire five-minute CHP test that followed incubation.

To further test of the reliability of CHP sensitivity, we included a “rekindling” phase (CHP rekindle; Dirks et al., 2003). After the initial CHP testing was completed, an icepack (see below for details) was applied to the forearm until a complete termination of pain was reported. Afterwards, the thermode was again placed over top of the site of capsaicin application, heated to 40°C, and held in place for five minutes. CHP rekindle scores were calculated as the average of the pain ratings provided during this five-minute period.

### Icepack Application

At the conclusion of the PHP and CHP tests, the thermode was removed and a disposable icepack was applied the stimulated area of the left forearm. This was done to prevent pain carryover from one test to another and to ensure that pain ratings for subsequent tests were captured from a starting state of no ongoing pain. The icepack was left in place until the complete absence of pain was reported by the participant. No participants indicated that the icepack itself was ever painful. Following each icepack application, a 5-minute pain-free, eyes closed EEG session occurred.

### Procedure

An outline of the experimental timeline and procedures is presented in Figure 1. In order to allow sufficient time for any long-term effects of capsaicin exposure to subside, visits were separated by 21 days or more (except for one case where a subject returned at 19 days because of a scheduling conflict; mean separation of Visit 1 and Visit 2 = 54.74 days, S.D. = 55.92 days, range = 19 – 310 days, Figure S1).

**Figure 1.**
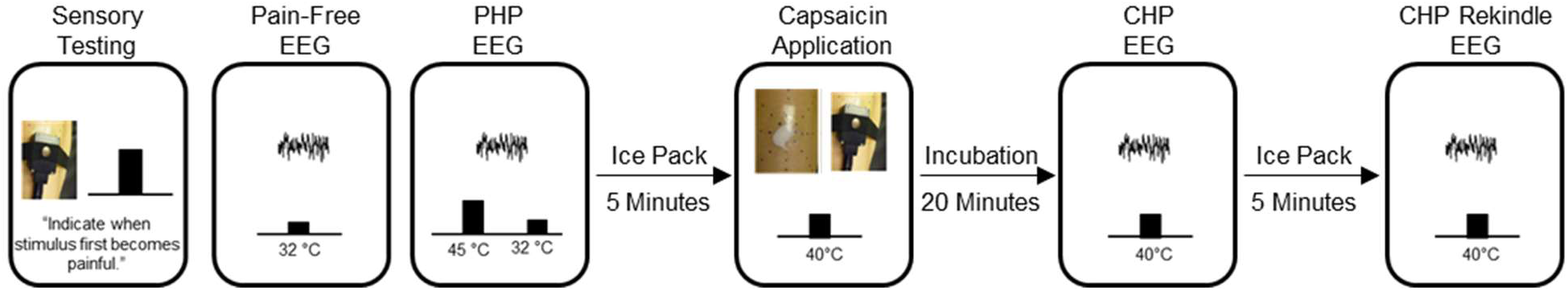
Outline of the experimental procedure. After a brief sensory testing session, participants completed a pain-free EEG. Next, participants completed a “Phasic Heat Pain (PHP)” EEG. After a 5-minute period in which an ice pack was applied to the skin, a second pain-free EEG was collected (not shown). Afterwards, capsaicin was applied to the forearm and incubated for twenty minutes. Next, a “Capsaicin-Heat Pain (CHP)” EEG was completed while a 40°C thermode was placed on top of the capsaicin. After a 5-minute period in which an ice pack was applied to the skin and a third pain-free EEG was collected (not shown), the 40°C thermode was again placed on top of the capsaicin and a 5-minute eyes-closed “CHP rekindle” EEG was completed. For each EEG, data was collected for 5 minutes while participants were instructed to keep their eyes closed. Identical procedures were performed at Visit 1 and Visit 2.

Participants first underwent an initial screening visit, Visit 0, that included quantitative sensory testing as well as additional tests to ensure that 40°C was rated as minimally-painful, to identify the appropriate PHP temperature, and to provide initial exposure to capsaicin. For the first four participants, these procedures, excluding capsaicin exposure, were performed during Visit 1.

Participants returned for Visit 1 at least three weeks after completing Visit 0. Most participants then returned at least three weeks after Visit 1 for Visit 2. Procedures for Visits 1 and 2 were identical. For the entirety of Visits 1 and 2, participants were seated in a comfortable chair in a quiet room that was isolated from strong electrical interference. For all EEG sessions, lights in the testing room were turned off and participants were instructed to close their eyes, remain still, relax without falling asleep, and continuously rate any pain they experienced with the manual analog scale placed at their right hand. Visits 1 and 2 began with quantitative sensory testing. For the first four participants, this sensory testing was not performed at Visit 2. After quantitative sensory testing, a brief 2-minute EEG was collected to ensure the quality of EEG recording. Next, a room temperature thermode was placed onto the left forearm while eyes closed, pain-free EEG was collected for 5 minutes. The primary objective of the current study was to use PAF recorded during this pain-free period as a predictor of subsequent pain sensitivity during CHP and PHP.

Following the pain-free EEG, prolonged pain was induced with the PHP model. During the five minutes of PHP, EEG was collected while participants rested with their eyes closed and continuously rated the intensity of any perceived pain. Upon completion of the PHP model, a disposable ice pack was placed onto the participant’s left forearm until they reported being completely free of pain after which 5 minutes of eyes closed EEG was collected. Next, the second model of prolonged pain, CHP, was induced. Participants were instructed to continuously rate the intensity of experienced pain during this incubation period.

Following the 20-minute incubation period, and with the thermode temperature still held at 40°C, 5 minutes of eyes closed, continuous EEG was recorded while participants continuously rated the intensity of any perceived pain. An icepack was then applied to the forearm and, once pain was reported to be completely absent, 5 minutes of eyes closed EEG was collected. Afterwards, a 40°C thermode was placed over the site of capsaicin application to induce CHP rekindling. Five minutes of eyes closed EEG was then recorded while participants continuously rated the intensity of any perceived pain.

### Data Processing

Because our primary objective was predicting pain sensitivity, the EEG data of interest were the initial pain-free EEGs collected at the beginnings of Visits 1 and 2. EEG data were preprocessed with EEGLAB 13.6.5b (Delorme and Makeig, 2004). Preprocessing began with filtering the data between .2 and 100Hz using a linear FIR filter. Channel data were then visually inspected and overtly noisy channels were removed from further analysis. Removed channels were not interpolated. On average, 1.64 (S.D. = 1.92, range: 0 – 8) and 1.79 (S.D. = 1.79, range: 0 – 6) channels were removed per individual from Visit 1 and Visit 2 datasets, respectively. Finally, Principal Components Analysis (PCA) was performed and components with spatial topographies and time series resembling blinks and/or saccades were removed from the data.

As opposed to our previous studies which used ICA to isolate alpha sources over visual and somatosensory regions, we used channel level data to increase the ease with which our methods can be reproduced. Although it may decrease the signal to noise ratio of the data, this approach eliminates the need to identify ICA components on a participant by participant basis and is equally effective for capturing the PAF-pain sensitivity relationship (Furman et al., 2019). For channel level analyses, we focused on channels (C3, Cz, and C4) that most strongly reflected the sensorimotor component topography observed in our original study (Furman et al., 2018). If a channel from this sensorimotor region of interest (ROI) was removed due to noise, only the remaining channels were used; this affected few participants (Visit 1: n = 4; Visit 2: n = 1) and no participant had more than one channel removed. In order to make the current results easily comparable to previous findings, all main analyses use PAF calculated from this sensorimotor ROI; use of this ROI is not intended to imply a mechanism or source for any documented effects.

To explore if additional EEG channels capture the PAF-pain sensitivity relationship, the surface Laplacian was computed following preprocessing (Perrin et al., 1989). Results from analyses using this estimate of current source density can be found in the Supplemental Data (Figure S6).

### Quantification of Sensorimotor PAF

The frequency decomposition of the sensorimotor ROI data was performed using routines in FieldTrip (Oostenveld et al., 2011). Data from each pain-free EEG session was segmented into non-overlapping 5 second epochs and power spectral density in the .2–100 Hz range (0.2 Hz bins) was derived with the ‘ft_freqanalysis_mtmfft’ function. A Hanning taper was applied to the data prior to calculating the spectra to reduce edge artifacts (e.g. Mazaheri et al., 2014).

At every channel and for each epoch, PAF was estimated using a center of gravity (CoG) method (Klimesch et al., 1993). We defined CoG as follows:

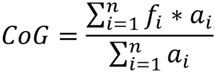

where *f_i_* is the ith frequency bin including and above 9 Hz, n is the number of frequency bins between 9 and 11 Hz, and *a_i_*-the spectral amplitude for *f_i_*. From our previous work, we have determined that this restricted frequency range reduces the influence of 1/f EEG noise on PAF estimation (Furman et al., 2018). Epoch-level PAF estimates were averaged to yield a single mean PAF estimate for each channel. Channel-level PAF estimates were further averaged across sensorimotor channels to yield a single sensorimotor PAF estimate for each participant at each visit.

To ensure that results were not an artifact of the range used for PAF estimation, PAF was additionally calculated with the wider 8-12 Hz range. Results with this wider estimation range are presented throughout the text and PAF estimates calculated using either the 9-11 or 8-12 ranges were highly similar (Figure S2).

### Statistical Analysis

All analyses were performed using custom scripts implemented in the Matlab environment (version R2013A). Statistical tests were conducted in Matlab or SPSS (Version 25).

Previous work has found that CHP evokes limited pain or hypersensitivity in roughly one third of individuals (Liu et al., 1998, Walls et al., 2017). While the reasons for this remain unclear, certain physiological factors, such as genetic polymorphisms (Campbell et al., 2009), appear to play a role in limiting the effects of the TRPV1 agonist itself. For this reason, it is difficult to determine whether insensitivity to capsaicin reflects a failure of the CHP model or an individual’s sensitivity to pain. To address this problem, we separated participants into three pain response classes: 1) individuals who display a clear pain response to CHP (average pain >= 10) at either Visit 1 or Visit 2 (“CHP responder”); 2) individuals who display a clear pain response to PHP at either Visit 1 or Visit 2 but no response to CHP at either visit (average pain < 10; “CHP non-responder”); and 3) individuals who do not display a clear pain response to PHP or CHP at either visit (“high tolerance”). For the high tolerance pain class, the presence of PHP insensitivity provides important evidence that CHP insensitivity is unlikely to reflect model failure alone. To ensure that results were not confounded by variability associated with an individual’s physiological ability to experience CHP, we chose to focus our main analyses on CHP responder and high tolerance individuals. For all tests involving PHP, results when including all three pain classes are also provided.

To determine if sensitivity to prolonged pain is similar across prolonged pain models, a series of pairwise correlations was calculated between all possible test pairs at each visit. For these and all other correlational analyses, Spearman’s rank order correlations were computed, and outliers were defined as data points greater than 2.5 standard deviations above or below the mean value obtained from Visit 1 data. We further assessed whether sensitivity is reliable across prolonged pain tests using Cronbach’s α.

To begin testing whether pain-free, sensorimotor PAF is related to prolonged pain sensitivity, we performed a series of pairwise correlations between pain-free, sensorimotor PAF and each pain test (PHP, CHP, and CHP rekindle) at each visit. Bonferonni corrections for multiple tests were applied to data from each visit (3 tests; one per prolonged pain test) yielding a corrected significance threshold of p = .017. For each test, we also investigated the effect of sex by performing correlations separately for males and females. To ensure that our results were not an artifact of our PAF estimation algorithm, we correlated pain sensitivity scores to the average, pain-free estimate of spectral power at each 0.2 Hz element within the 8-12 Hz range. For this analysis, spectra were z-scored in order to normalize total spectral power across individuals.

Next, we determined whether pain-free, sensorimotor PAF can accurately identify the most or least pain sensitive individuals. In the first analysis, we used a series of linear support vector machines (SVM) to perform leave-one-out, within-study classification (internal validation). To do so, pain scores from PHP, CHP, and CHP rekindle were averaged and, in separate tests, the top or bottom 10% of averaged pain scores were labelled as targets. A series of SVMs were then trained to identify targets based on Visit 1 baseline, pain-free PAF estimates from all but one individual (training set). Trained support vector machines were then used to predict whether the withheld participant was a target. Visit 1 data was used in order to maximize the size of the available dataset. Each participant served as the test exactly once and predictions were evaluated using F_1_ scores (harmonic mean of precision and recall; Sokolova & Lapalme, 2009; Lipton et al., 2014). F_1_ scores are often used when the proportions of two classes are uneven. To determine the full scope of prediction, we repeated this analysis by increasing the percentage of data labelled as a target in increments of 10% up to a maximum of 50% (i.e. median split of data). To evaluate F_1_ scores, we generated a distribution of null F_1_ scores by assigning targets at random and then performing the analysis described above. This procedure was carried out 10,000 times and obtained F_1_ scores were evaluated as significant if they were equal to or surpassed the 95^th^ percentile of the null distribution.

In the second analysis, we used a single linear SVM to perform cross-study classification using data from the current study as the training set and data from an earlier study on CHP sensitivity as the test set (external validation; Furman et al., 2018). Prior to analysis, PAF estimates within each study were normalized to z-scores. Otherwise, details of this analysis were identical to those of the within-study classification analysis.

To examine whether pain-free, sensorimotor PAF is reliable across Visits 1 and 2, estimates from each visit were compared using a paired *t*-test. Bayes factor analysis was used to determine whether the null hypothesis could be accepted (i.e. no change in PAF between visits). Bayes factor analysis provides a method for assessing the relative evidence in favor of either the null or alternative hypothesis with a Bayes factor less than .33 or greater than 3 are taken as strong evidence in favor of the null and alternative hypotheses, respectively (Rouder et al., 2009); Bayes factor scores in-between these values are considered to provide no evidence in favor of either hypothesis. As an additional test of stability, PAF estimates at Visits 1 and 2 were correlated with one another.

The stability of prolonged pain scores was assessed using a linear mixed effects model with subjects as random effects (intercept included) and Visit (Visit 1 vs Visit 2), Type (WDT vs. HPT vs. Phasic vs. CHP), and the Visit X Type interaction as fixed effects. We were specifically interested in determining whether scores change over time (main effect of Visit) and whether these changes were specific to individual tests (Visit X Type interaction). For each prolonged pain test, Bayes factor analysis was used to determine whether the null hypothesis could be accepted (i.e. no change in pain score between visits). Additionally, the stability of pain scores from each test was analyzed by correlating Visit 1 and Visit 2 pain scores.

To further test of the stability of pain-free, sensorimotor PAF and prolonged pain scores, we examined the correlation between pain-free, sensorimotor PAF at Visit 1 and Visit 2 pain sensitivity. To ensure that results were not an artifact of our PAF estimation algorithm, we also correlated pain sensitivity scores to the average, pain-free estimate of spectral power at each 0.2 Hz element within the 8-12 Hz range. Finally, we tested whether pain-free, sensorimotor PAF at Visit 1 could accurately identify the least and most pain sensitive individuals at Visit 2. As before, a series of leave-one-out SVMs were trained to identify the least or most pain sensitive individuals and then tested on the withheld participant. Performance was quantified by comparing the observed F_1_ score to a bootstrapped, null distribution of F_1_ scores.

## Results

From our initial cohort of 61 individuals, two individuals were removed due to abnormal pain ratings: one participant fell asleep during ratings while another participant provided extremely high pain ratings in the absence of any noxious stimuli indicating that they may been confused by the rating scheme. We excluded one additional participant who experienced a change in CHP score, + 69.26, that was 3.82 standard deviations greater than the average CHP change (average change = 1.76, S.D. = 17.64). No other change in CHP scores was greater than 2.05 standard deviations above the mean (range = +37.96 to −31.05).

From the remaining 58 participants (Table 1), 33 participants were classified as CHP responders (CHP score > 10), 10 participants were classified as high tolerance individuals (CHP and PHP scores < 10), and 15 participants were classified as CHP non-responders (PHP score > 10 & CHP score < 10). Due to a technical error, EEG data was lost for one CHP responder at Visit 1; Visit 1 data for this individual was only included in prolonged pain analyses. Of the 58 individuals providing data at Visit 1, a total of 43 individuals also provided data at Visit 2, of which 32 had been classified as a CHP responder or high tolerance individual. CHP rekindle data for one participant at Visit 2 was not collected. Unless otherwise stated, analyses only include data from high tolerance and CHP responder individuals.

A summary of prolonged pain scores for each pain response classification is presented in Figure 2. Both PHP and CHP produced sensitization, a hallmark of prolonged pain (see Supplemental Data), and similar amounts of pain in males and females (Supplemental Figure S3). Correlations between all possible pairs of tests were significant (Table 2) and this conclusion held when analyses were repeated while including all participants regardless of pain response classification (Supplemental Figure S1B). Reliability analysis further revealed that sensitivity was consistent across prolonged pain tests, Chronbach’s α = .91 (Visit 1 alone, α = .82, Visit 2 alone, α = .83). Including all subjects, regardless of pain response classification, did not alter this finding, Chronbach’s α = .90 (Visit 1 alone, α = .77, Visit 2 alone, α = .83). Thus, CHP and PHP appear to sample similar prolonged pain processes.

**Figure 2.**
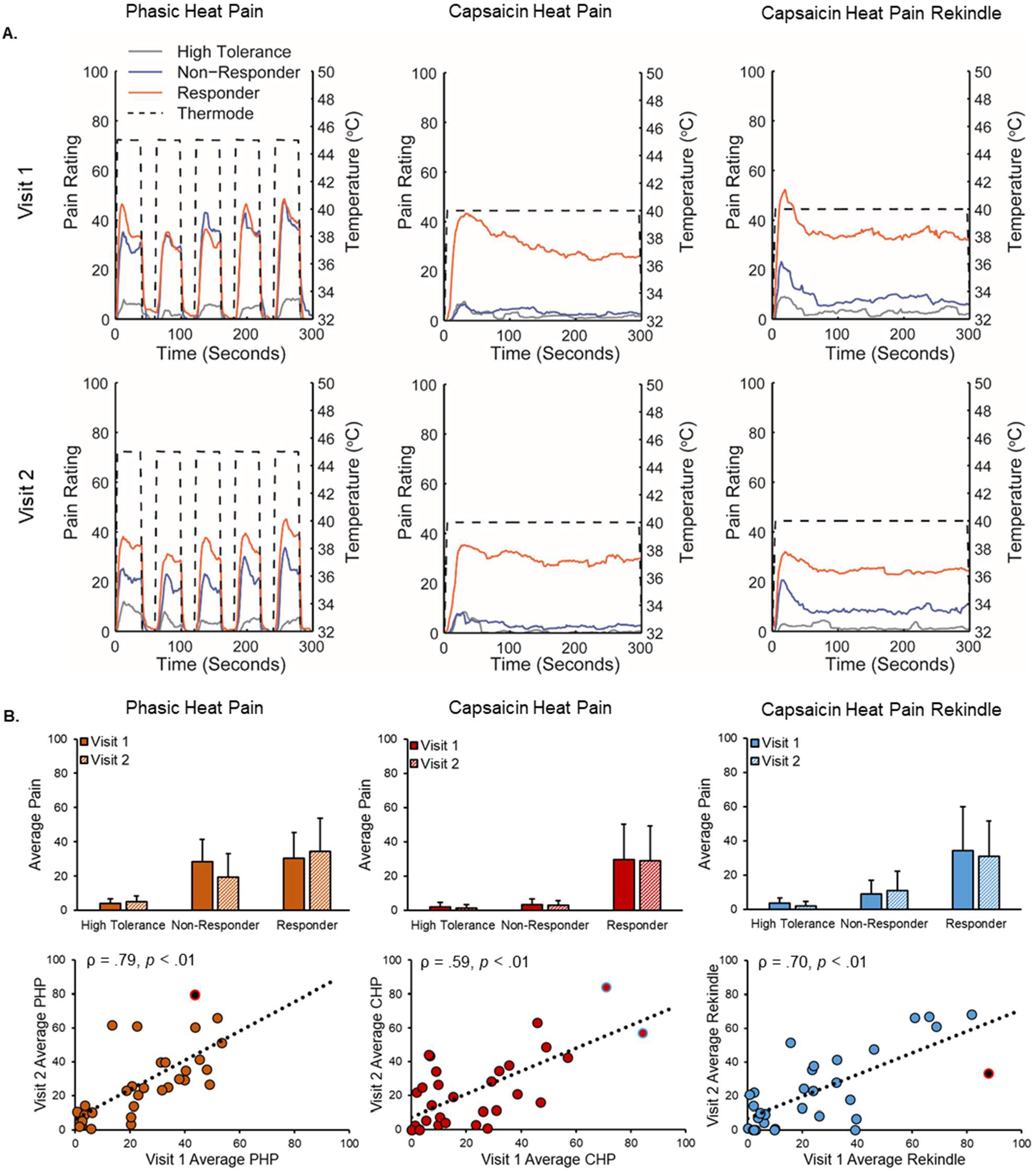
Prolonged pain models were stable across visits. A. Average pain time courses for CHP responder (orange), CHP non-responder (blue), and high tolerance (grey) pain classifications during each prolonged pain test. Dotted lines reflect the temperature of the thermode applied to the forearm during each test. CHP responders were defined as participants with pain scores >10/100 in response to CHP, CHP non-responders were defined as participants who had pain scores <10/100 during CHP and pain scores >10/100 during PHP, and high pain tolerance individuals were defined as participants with pain scores <10/100 in response to both CHP or PHP. B. Pain ratings broken down by prolonged pain test, pain response classification, and visit. Bar graphs reflect means and error bars reflect +1 standard deviation. Scatter plots only include data from CHP responders and high tolerance individuals. Off-color data points represent statistical outliers not included in analyses and dotted lines represent the linear regression line of best fit.

**Table 2.** Spearman correlation coefficients (*p* values) between sensory tests at each testing visit.

|  |  | Visit 1 |  |  | Visit 2 |  |  |
| --- | --- | --- | --- | --- | --- | --- | --- |
|  |  | PHP | CHP | CHP Rekindle | CHP Rekindle | CHP | PHP |
| PHP |  |  | .52 (<.016) | .50 (<.016) | .58 (<.016) | .77 (<.016) |  |
| CHP |  |  |  | .88 (<.016) | .78 (<.016) |  |  |
| CHP Rekindle |  |  |  |  |  |  |  |

### Sensorimotor PAF is Reliably Predicts Thermal, Prolonged Pain Sensitivity

At Visit 1, pain-free, sensorimotor PAF predicted pain sensitivity to all three prolonged pain tests, PHP: Spearman ρ = −.43, p < .01; CHP: Spearman ρ = −.44, p < .01; CHP rekindle sensitivity, Spearman ρ = −.44, p < .01 (Figure 4). Similar results were obtained for PHP when we used a partial correlation to account for variability in the thermode temperature used during PHP, Spearman *ρ* = −.40, *p* = .01, or when we included all participants regardless of pain response classification, Spearman *ρ* = −.34, *p* = .01. Expanding the PAF calculation range to 8 – 12 Hz did not greatly impact the relationship for any prolonged pain test, PHP: Spearman ρ = −.38, *p* = .01; CHP: Spearman ρ = −.34, *p* = .03, CHP rekindle: −.38, *p* = .01 (Supplemental Figure S2E). Furthermore, inspection of the relationship between pain sensitivity and power at each frequency element within the alpha range demonstrate that these results are not an artifact of our PAF calculation method: for each test, slower (8-9.5 Hz) elements were positively associated with pain sensitivity while faster (10.5-12 Hz) elements were negatively associated with pain sensitivity (Figure 4 Lower Panels). We found no evidence of sex effects on the relationship of PAF to either PHP, CHP, or CHP rekindle (Supplemental Figure S4A). Interestingly, the relationship between PAF and prolonged pain sensitivity was apparent at nearly every scalp channel even when volume conduction was accounted for with a surface Laplacian transformation (Supplemental Figures S5 & S6).

**Figure 3.**
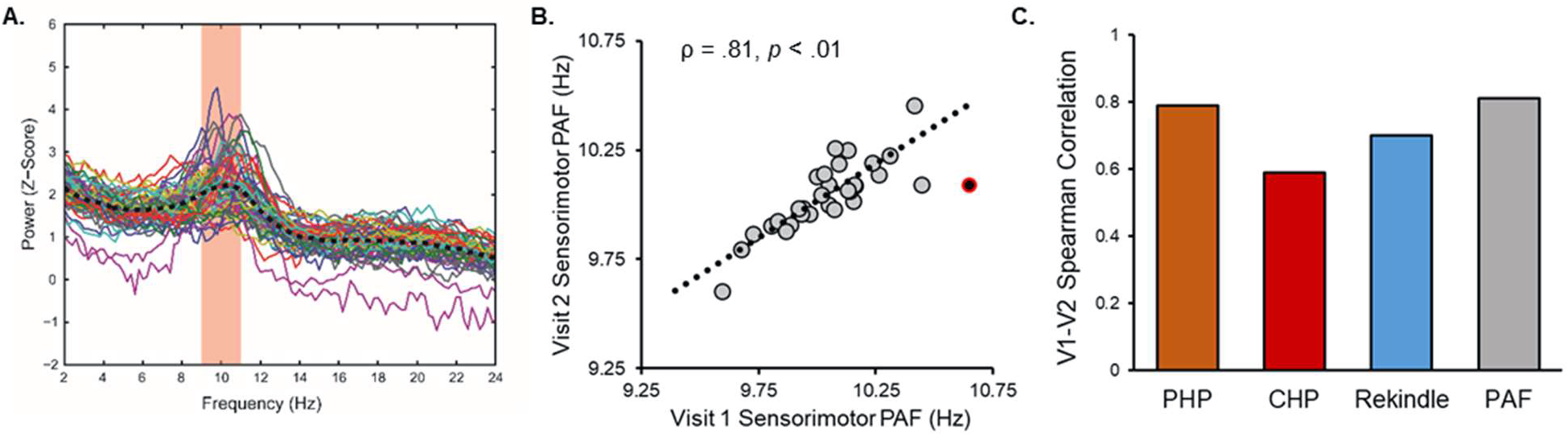
A. Pain-free, sensorimotor ROI spectra collected from all participants Visit 1. Colored lines reflect individual participants, and the black dashed line reflects the average spectra across all participants. The red zone reflects the frequency range (9-11 Hz) used to calculate PAF according to the center of gravity method. B. Pain-free, sensorimotor PAF estimates are strongly correlated across Visits. Note that Visit 2 occurred, on average, 7.8 weeks after Visit 1. Off-color data points represent statistical outliers not included in analyses and the dotted line represents the linear regression line of best fit. C. Pain-free, sensorimotor PAF and prolonged pain models are stable across Visits.

**Figure 4.**
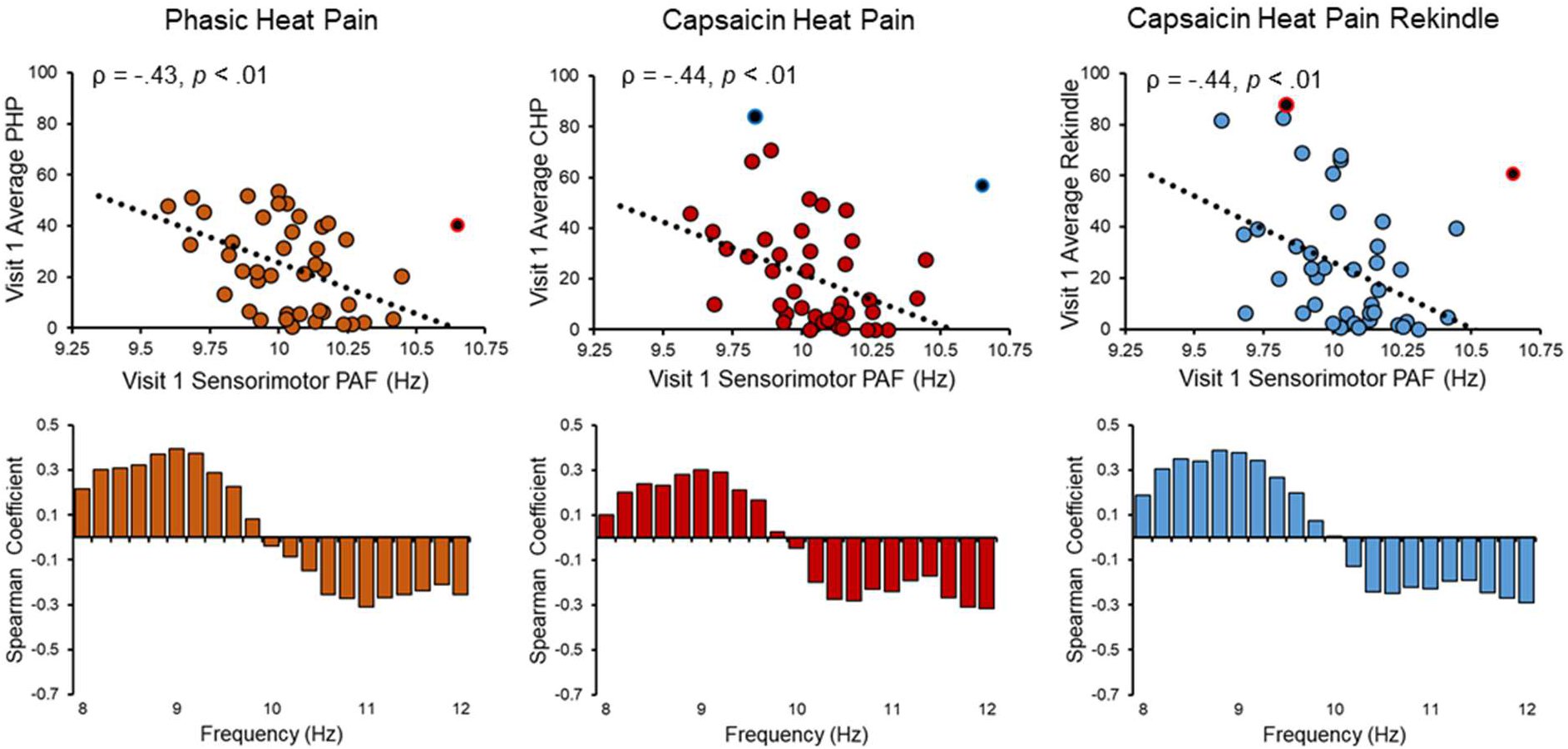
Visit 1 pain-free, sensorimotor PAF is correlated with sensitivity to all three Visit 1 prolonged pain tests. Off-color data points represent statistical outliers not included in analyses and dotted lines represent the linear regression line of best fit. Bar graphs below each scatter plot reflect Spearman correlation coefficients between Visit 1 pain scores and Visit 1 estimates of pain-free power at each 0.2 Hz bin within the 8-12 Hz range. For all three tests, frequency elements below 10 Hz are positively associated with pain sensitivity while frequency elements above 10 Hz are negatively associated with pain sensitivity.

At Visit 2, pain-free, sensorimotor PAF again predicted pain sensitivity to all three prolonged pain tests, PHP: Spearman ρ = −.59, p < .01; CHP: Spearman ρ = −.57, p < .01; CHP rekindle sensitivity, Spearman ρ = −.43, p = .016 (Figure 5). As before, PHP outcomes remained stable when either accounting for thermode temperature with a partial correlation, Spearman *ρ* = −.55, *p* < .01, or including all 43 participants regardless of pain response classification, Spearman *ρ* = −.37, *p* = .02. Expanding the PAF calculation range to 8-12 Hz did not impact PAF’s relationship to any test, PHP: Spearman ρ = −.51, p < .01; CHP: Spearman ρ = −.58, p < .01, CHP rekindle: Spearman ρ = −.44, p = .01 (Supplemental Figure S2E), and correlations between pain and power across the alpha range once again revealed an association of slow and fast ranges with heightened and decreased pain sensitivity, respectively (Figure 5 Lower Panels). As in Visit 1, there did not appear to be an influence of sex on the relationship between PAF and any of our prolonged pain tests (Figure S4A) and this relationship was evident across the entire scalp (Figures S5 & S6).

**Figure 5.**
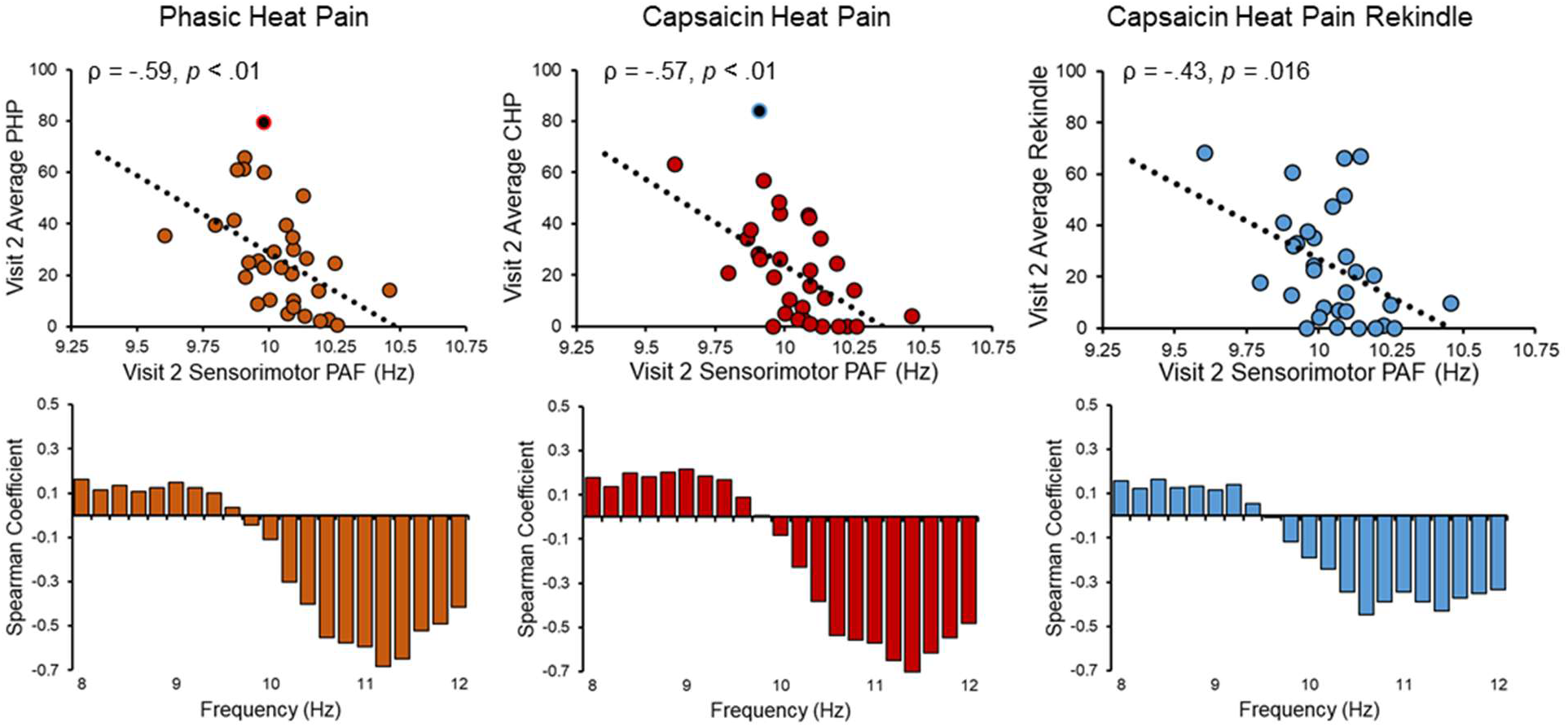
Visit 2 pain-free, sensorimotor PAF is significantly correlated with sensitivity to all three Visit 2 prolonged pain tests. Off-color data points represent statistical outliers not included in analyses and dotted lines represent the linear regression line of best fit. Bar graphs below each scatter plot reflect Spearman correlation coefficients between Visit 2 pain scores and Visit 2 estimates of pain-free power at each 0.2 Hz bin within the 8-12 Hz range. For all three tests, frequency elements below 10 Hz are positively associated with pain sensitivity while frequency elements above 10 Hz are negatively associated with pain sensitivity.

### Sensorimotor PAF Can Identify the Most Pain Sensitive Individuals

Given the high sensitivity of classification analyses to outliers, one participant with an extreme PAF estimate was not included in either analysis (PAF = 10.65, 3.20 S.D. above the mean). In order to make the classification analysis generalizable to other datasets, and to take advantage of the strong correlation between prolonged pain tests, we created a composite pain sensitivity score by averaging scores from all three prolonged pain tests (PHP, CHP, CHP Rekindle). This pain sensitivity score was significantly correlated with PAF at both Visit 1, Spearman ρ = −.51, p < .01, and Visit 2, Spearman ρ = − .60, p < .01 (Supplemental Figure S7A). This relationship remained evident when we included all participants regardless of classification, Visit 1: Spearman ρ = −.42, p < .01; Visit 2: Spearman ρ = −.33, p = .03 (Supplemental Figure S7A).

Support vector machines (SVM) trained and tested on the current dataset were able to identify both the least and most sensitive individuals using just pain-free PAF estimates (internal validation; details found in the Statistics section). Compared to a simulated null distribution of F_1_ scores, the least pain sensitive individuals were identified at above chance levels at all labelling intervals but the 20% one (Figure 7B). Similarly, the most sensitive individuals were identified at above chance levels at all labelling intervals but the 30% one. When including all participants, regardless of classification, PAF significantly identified the least sensitive individuals at all labelling intervals but only the most sensitive individuals at the 10% and 50% intervals (Supplemental Figure S7B). This latter result likely reflects that the composite pain sensitivity score fails to capture the mixed sensitivity of CHP non-responders to CHP and PHP.

A linear SVM trained on the current dataset could identify high and low pain sensitive individuals in a separate, independent study (external validation). Using a similar procedure to one used for within-study classification, a single linear SVM trained on data from the current study was used to predict the identity of 21 participants from a previous study on CHP sensitivity (Furman et al., 2018). Compared to a simulated null distribution of F_1_ scores, we found that PAF estimates identified the most pain sensitive individuals at above chance levels for all labelling intervals and identified the least pain sensitive individuals at above chance levels only at the two largest, 40% & 50%, intervals (Figure 7D). Rerunning the analysis with all participants, regardless of pain response classification, included in the training set yielded identical results (Supplemental Figure S7B)

### Sensorimotor PAF and Prolonged Pain Sensitivity Are Stable over Time

One possible explanation for the presence of a reliable relationship between pain-free, sensorimotor PAF and prolonged pain sensitivity at Visits 1 and 2 is that both measures are themselves stable over time. In line with this premise, Visit 1 (mean = 10.04, S.D. = .20) and Visit 2 (mean = 10.04, S.D. = .16) estimates of pain-free, sensorimotor PAF were not significantly different, *t*(29) = .32, p = .75, and Bayes factor analysis supported the null hypothesis of no differences between the two, Bayes Factor < .01. These results did not change when we included all participants regardless of pain response classification *t*(40) = .34, p = .73, Bayes Factor < .01. What’s more, Visit 1 and Visit 2 estimates of pain-free, sensorimotor PAF were strongly correlated, Spearman ρ = .81, p < .01 (Figure 3A); this finding did not change when we including all participants regardless of pain response, Spearman ρ = .82, p < .01, or expanded the PAF calculation range to 8-12 Hz, Spearman ρ = .86, p < .01.

Similarly, a linear mixed effects model revealed that prolonged pain sensitivity did not change over time with neither the main effect of Visit, *F*_(1,161.32)_ = .13, *p* = .72, nor the Visit x Pain Type interaction, *F*_(2,113.244)_ = .26, *p* = .77, reaching significance. Bayes factor analysis failed, however, to support either the null or alternative hypothesis for any prolonged pain test, PHP: Bayes Factor = 1. 19, CHP: Bayes Factor = .71, CHP Rekindle: Bayes Factor = .74. Visit 1 and Visit 2 pain scores were correlated for all three prolonged pain tests, PHP, ρ = .79, p < .01, CHP, ρ = .59, p < .01, and CHP rekindle, ρ = .70, p < .01 (Figure 2), and remained so when we expanded the dataset to include CHP non-responders, PHP: ρ = .74, p < .01; CHP: ρ = .69, p < .01, CHP Rekindle, ρ = .68, p <. 01.

### Sensorimotor PAF Can Predict Thermal, Prolonged Pain Sensitivity Occurring 8 Weeks Later

If pain-free, sensorimotor PAF and prolonged pain sensitivity are stable traits then Visit 1 PAF should be able to predict Visit 2 pain scores collected, on average, 8 weeks later. Indeed, we found that Visit 1 pain-free, sensorimotor PAF and Visit 2 pain scores were strongly correlated, PHP: Spearman *ρ* = −.67, *p* < .01; CHP: Spearman *ρ* = −.62, *p* < .01; CHP rekindle: Spearman *ρ* = −.52, *p* < .01 (Figure 6). For PHP, this relationship remained when we controlled for variability in thermode temperature, Spearman *ρ* = −.66, *p* < .01, or included all participants regardless of pain response classification, Spearman *ρ* = −.44, *p* < .01.

**Figure 6.**
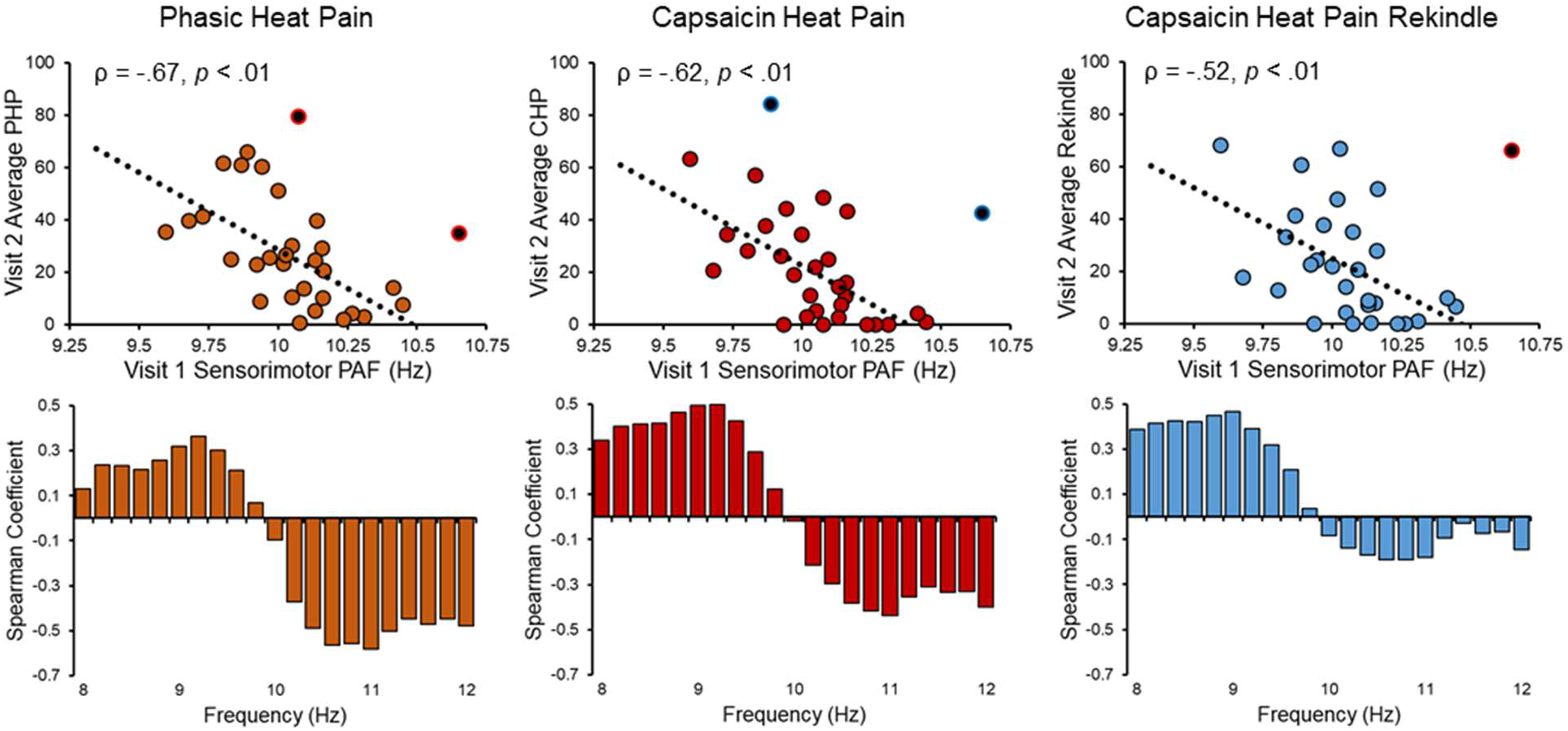
Visit 1 pain-free, sensorimotor PAF predicts sensitivity to all three Visit 2 prolonged pain tests. Note that Visit 2 occurred, on average, 7.8 weeks after Visit 1. Off-color data points represent statistical outliers not included in analyses and dotted lines represent the linear regression line of best fit. Bar graphs below each scatter plot reflect Spearman correlation coefficients between Visit 2 pain scores and Visit 1 estimates of pain-free power at each 0.2 Hz bin within the 8-12 Hz range. For all three tests, frequency elements below 10 Hz are positively associated with pain sensitivity while frequency elements above 10 Hz are negatively associated with pain sensitivity.

Expanding the PAF calculation range to 8-12 Hz did not impact PAF’s relationship to any test, PHP: Spearman ρ = −.57, p < .01; CHP: Spearman ρ = −.52, p < .01, CHP rekindle: Spearman ρ = −.45, p = .015, and correlations between pain and power across the alpha range again demonstrated that the slow and fast ranges were associated with heightened and decreased pain sensitivity, respectively (Figure 6 Lower Panels).

What’s more, the least and most pain sensitive individuals at Visit 2 could be identified using Visit 1 pain-free, sensorimotor PAF. Visit 2 pain sensitivity, represented as the average pain score across tests, was strongly correlated to Visit 1 pain-free, sensorimotor PAF, Spearman *ρ* = −.66, *p* < .01, and remained so when including all participants regardless of pain classification, Spearman *ρ* = −.45, *p* < .01 (Supplemental Figure S7A). Compared to a null distribution of F_1_ scores, pain-free, sensorimotor PAF identified the most sensitive individuals at all but the smallest labelling interval and the least sensitive individuals at all but the two smallest labelling intervals (Figure 7E). Classification failure at the smallest labelling intervals was likely due to the relatively low number of targets available (sample = 30; targets = 3 and targets = 6 at the 10% and 20% labelling intervals, respectively). Rerunning the analysis with all participants, regardless of pain response classification, again demonstrated that pain-free, sensorimotor PAF could identify the most sensitive individuals at all labelling intervals but the smallest one. For the least pain sensitive individuals, pain-free, sensorimotor PAF failed to yield significant predictions at any labelling interval (Supplemental Figure S7B).

**Figure 7.**
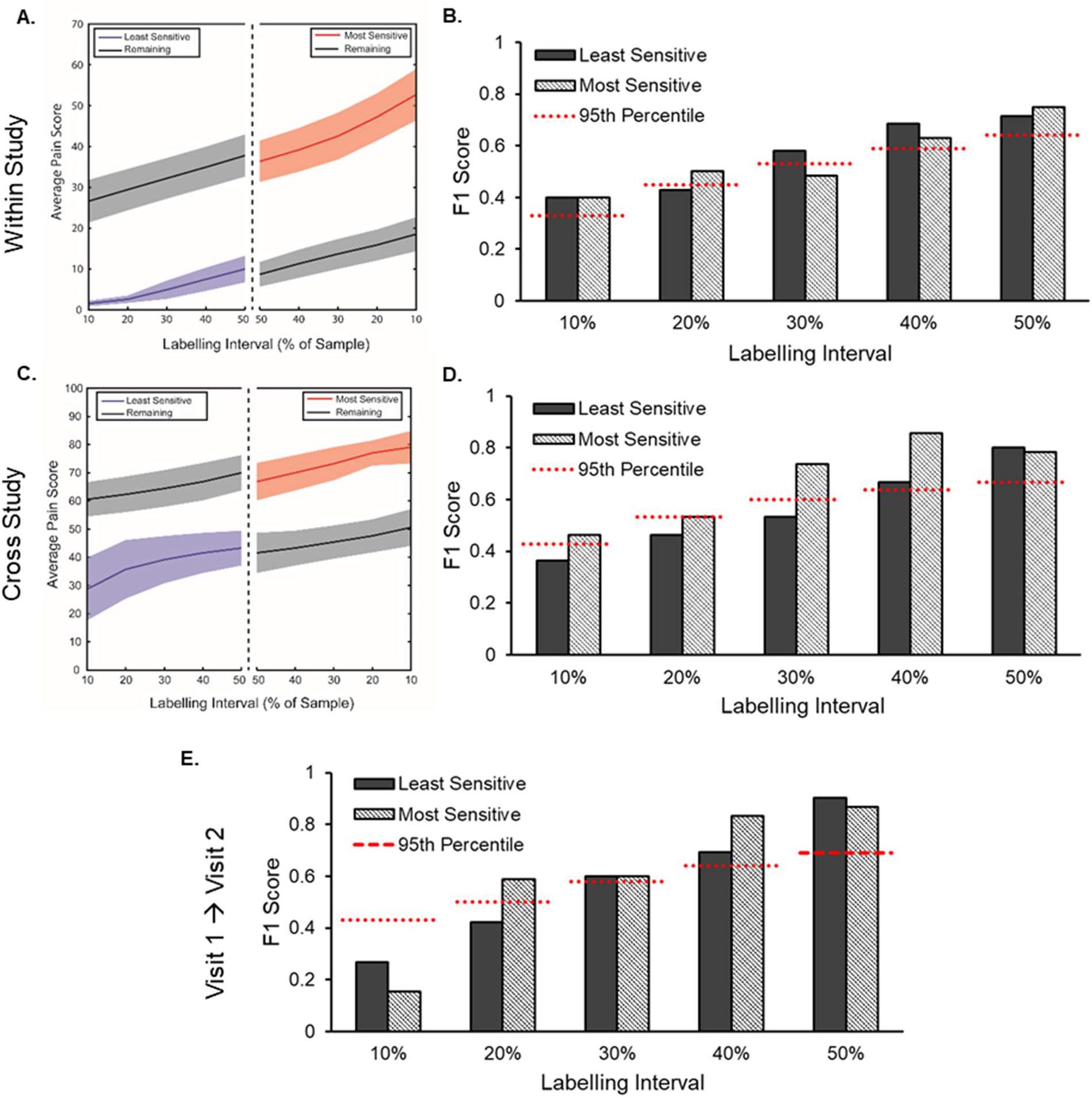
Visit 1 pain-free, sensorimotor PAF can accurately predict the identity of the most pain sensitive individuals and a support vector machine trained on this data can identify the most pain sensitive in an independent study. A. Visit 1 pain scores from the three prolonged pain models were averaged and a relevant percentage of the sample, ranging from 10% to 50% (i.e. median split), was identified as high or low pain sensitive. Colored lines (shading = 95% confidence interval) reflect the average sensitivity for identified participants and those not classified (black lines; “Remaining”). B. A support vector machine trained on Visit 1 pain-free, sensorimotor PAF predicts the identity of high and low pain sensitive individuals from the same study at almost all labelling intervals. An F_1_ score of 1 indicates perfect classifier performance and the dashed red lines reflect the 95^th^ % of a null distribution of F_1_ scores. C. Same as in A., except pain scores were taken from an independent study on PAF and CHP (Furman et al., 2018). D. A support vector machine trained on Visit 1 pain-free, sensorimotor PAF predicts the identity of high pain sensitive individuals from an independent study at all labelling intervals. E. A support vector machine trained on Visit 1 pain-free, sensorimotor PAF predicts the identity of Visit 2 high pain sensitive individuals. Note that pain scores for this test are not provided but are nearly identical to those present in C.

## Discussion

Cycles of the 8 – 12 Hz Alpha oscillation are thought to reflect rhythmic, inhibitory processes that control the temporal dynamics of sensory processing (Jensen & Mazaheri, 2010; Van Rullen 2016). Peak Alpha Frequency (PAF), the individual-specific frequency at which these rhythms are dominantly expressed, is thought to reflect the speed at which sensory information is sampled (e.g. Samaha & Postle, 2015; Cecere et al., 2015; Wutz et al., 2018). PAF abnormalities are evident in several chronic pain conditions, with patients often demonstrating slowed PAFs relative to age-matched controls (e.g. Sarnthein et al., 2006; de Vries et al., 2013; Lim et al., 2016). These findings have led to proposals that PAF disturbances reflect ongoing, pathological processes such as Thalamocortical Dysrhythmia (e.g. Llinás et al., 1999). PAF, however, also appears to play a role in shaping the sensitivity of healthy individuals to prolonged pain (Nir et al., 2010; Furman et al., 2018; Furman et al., 2019). We have previously shown that the speed of PAF collected in the absence of a noxious stimulus is negatively related to an individual’s sensitivity to future prolonged pain events (i.e. slower PAF = greater pain sensitivity). This has led us to propose that pain-free PAF is a biomarker of prolonged pain sensitivity and, furthermore, that chronic-pain related disturbances of PAF may reflect differences in pain sensitivity that predate disease onset.

In the current study we examined the relationship of pain-free PAF to two models of prolonged pain, Capsaicin Heat Pain (CHP) and Phasic Heat Pain (PHP), within the same group of participants at two separate timepoints. From these experiments, we present two key pieces of evidence supporting the hypothesis that pain-free PAF is a prolonged pain sensitivity biomarker. First, pain-free PAF shares a near identical, negative relationship to CHP and PHP sensitivity, with increasingly slower PAF associated with increasingly greater pain intensity during each test. While we have previously reported a relationship between pain-free PAF and CHP sensitivity (Furman et al., 2018), the described relationship to PHP sensitivity is entirely novel. Reproduction of this relationship across models, despite differences in their length of application, the temperatures used, and the presence of a sensitizing agent, provides important evidence that PAF is a marker of prolonged pain sensitivity per se and not specific portions of either model. This interpretation is also supported by the replication of our earlier CHP findings despite large procedural differences between the two studies (i.e. CHP preceded by a cognitive or separate pain task). Preservation of the PAF-pain sensitivity relationship through the rekindling phase of the CHP model provides yet another piece of evidence that PAF captures an element of the prolonged pain experience that is independent of the local context (i.e. continuous vs. interrupted pain). Although the association of pain-free PAF with non-thermal forms of prolonged pain was not tested in the current study, similar findings in a musculoskeletal model of prolonged pain provide some assurance that pain-free PAF is likely to apply to a wide range of prolonged pain modalities (Furman et al., 2019).

Second, the relationship between pain-free, PAF and prolonged pain sensitivity is reliable over time. Within the same set of individuals, we show that the relationship between pain-free PAF and prolonged pain sensitivity is present at two separate testing visits. It should be acknowledged that this relationship was qualitatively stronger at Visit 2, which could be interpreted as evidence that factors that change with repeated testing, such as participant familiarity and/or vigilance, mediate the connection between pain-free PAF and prolonged pain sensitivity. While these effects cannot be entirely discounted, a separate explanation centers on the limited participant sample available at Visit 2; restricting Visit 1 analyses to only those participants completing both visits revealed relationship magnitudes, PHP: ρ = − .50; CHP: ρ = −.50; CHP rekindle: −.52, closer to those found at Visit 2.

This temporally stable association of pain-free PAF and prolonged pain sensitivity appears to be a consequence of the temporal stability of the measures themselves. For both pain-free PAF and prolonged pain sensitivity, we found that Visit 1 and Visit 2 estimates were strongly correlated and did not significantly differ from one another. These findings fit well with previous studies of PAF and prolonged pain sensitivity that demonstrate each are trait-like measures (e.g. Grandy et al., 2013; Naert et al., 2008; Koenig et al., 2014). Importantly, the average length of time separating visits (~ 8 weeks), as well as the absence of visual and haptic feedback during rating, provides comfort that the reliability of pain scores is not simply the result of participant’s explicit recollection of previous pain. From a broader perspective, these findings suggest the that relationship between pain-free PAF and prolonged pain is not uniquely determined at each visit but is instead an association that remains consistent *across* time; put another way, the *same* pain-free, PAF and the *same* prolonged pain sensitivity are sampled from individuals at each visit. Indeed, the ability of Visit 1 pain-free PAF to predict prolonged pain sensitivity at Visit 2 provides strong evidence in favor of this conclusion. Thus, these findings clearly show that pain-free PAF can provide cogent information about prolonged pain sensitivity at both short (i.e. minutes/hours separating PAF acquisition and pain testing) and long (weeks/months separating visits) timescales.

Considering its apparent reliability as a pain sensitivity biomarker, as well as its ease of obtainment, pain-free PAF holds real promise as a pain management and prophylaxis tool. This may be especially true in cases of planned surgery, where post-operative pain sensitivity is consistently found to be an important risk factor for chronic pain development (Hah et al., 2019). For example, identification of high pain sensitivity with PAF could be used to inform clinician decision making about surgical alternatives. To evaluate this possible real-world application, we examined whether pain-free PAF can predict the identify of high or low pain sensitive individuals. In almost all cases, a support vector machine trained on pain-free PAF was able predict the identity of the most pain sensitive individuals. This held true when the test data came from the current study or when it originated from an entirely separate study (Furman et al., 2018). In contrast, identification of the least pain sensitive individuals occurred when classification was applied to data from the current study but not when applied to outside data. These results suggest that pain-free PAF is particularly well suited for identifying high pain sensitive individuals. Importantly, Visit 1 pain-free PAF could be used to predict high pain sensitivity at Visit 2 suggesting that pain sensitivity prediction remains relevant across clinically relevant periods of time. Prospective collection of pain-free PAF at routine check-ups may therefore prove an effective strategy for ensuring information about an individual’s pain sensitivity is available to clinicians in cases of unplanned surgical intervention.

Despite the promise of the current findings, some potential limitations must be acknowledged. First, a subset of individuals demonstrating insensitivity to CHP were not included in the main set of analyses. Although a wide range of factors can render an individual less sensitive to capsaicin, at least some cases appear to be determined by physiological factors, like genetic polymorphisms (Campbell et al., 2009), that limit the effects of the TRPV1 agonist itself (i.e. model failure). The sources of pain sensitivity for these individuals and for those susceptible to the full range of capsaicin effects are thus fundamentally different and not comparable. This represents a limitation of the CHP model and not, in our opinion, a limitation of PAF’s ability to reflect pain sensitivity. To overcome this potential pitfall, we only included CHP-insensitive individuals if they also reported minimal pain in response to PHP. In these cases, the presence of PHP insensitivity provided important evidence that CHP insensitivity was at least partly attributable to an individual’s high tolerance of pain and not just model failure. While this decision could be interpreted as a confound to analyses of PHP, where sensitivity to capsaicin is not a relevant factor, supplementary results when all participants were included are provided for each test and, in all cases, conclusions regarding the relationship of pain-free PAF and PHP remained unchanged. Similarly, averaging pain scores across tests revealed that, even when including all participants, this broader description of pain sensitivity was well described by pain-free PAF. As a result, we feel confident that pain-free PAF’s relationship to pain sensitivity holds broadly across individuals.

Additionally, the current study is unable to provide concrete information about PAF’s source or identity. For the sole purpose of remaining consistent with our earlier methods, we chose to explicitly focus on PAF recorded from sensorimotor channels. As we have noted previously (Furman et al., 2019), PAF’s relationship to pain sensitivity is not restricted to sensorimotor channels and instead appears to encompass nearly every scalp channel. This continued to hold true in the current study even when possible volume conduction effects were controlled with a Laplacian transform. Although considered a limitation here, the widespread nature of PAF’s relationship to pain sensitivity may provide an important clue to its identity. In line with findings that the alpha rhythm travels across the cortex in “waves” (Zhang et al., 2018; Lozano-Soldevilla et al., 2019), PAF may reflect processes or sources whose actions are distributed across the brain. The thalamus represents one obvious candidate given its extensive cortical projections (e.g. Behrens et al., 2003) and central role in generating the alpha rhythm (Hughes and Crunelli, 2005). Large-scale, functional networks like those involved in attention also represent promising possibilities. Among these, the frontoparietal network is particularly interesting given that its relationship to the alpha rhythm is speed dependent (Sauseng et al., 2005; Sadaghiani et al., 2012) and has itself been implicated in individual differences in pain sensitivity (Kong et al., 2013; Tu et al., 2019). Resolution of this question will ultimately require both spatially sensitive methods, like EEG-fMRI, and careful behavioral testing to determine the brain regions and processes which mediate the relationship between pain-free PAF and pain sensitivity.

Some readers may also be concerned with the limited, 9-11 Hz frequency range that was used to calculate PAF. Alpha activity is not limited to 9-11 Hz range and has even been suggested to extend beyond the canonical 8-12 Hz range (Haegens et al., 2014). One advantage of the restricted calculation range we employed is that it most effectively negates the impact of the 1/*f* aperiodic signal on PAF estimation (Furman et al., 2018). While methods for isolating narrowband signal from aperiodic signal are advancing quickly (i.e. Haller et al., 2018), we found that they were unable to generate adequate solutions for all participants. As a result, we chose to focus on the 9-11 Hz range in order to provide the cleanest possible estimate of PAF. Importantly, results for all analyses were unchanged when PAF was calculated using the full 8-12 Hz range. Similarly, correlations of pain with estimates of spectral power at each 0.2 Hz element within the 8-12 Hz range confirmed that this relationship is not an artifact of either the range or method used to calculate PAF. Frequency elements below 10 Hz showed a consistent, positive relationship to pain sensitivity whereas elements above 10 Hz were negatively associated with pain. This finding reinforces that *where* power is expressed within the alpha range is relevant to pain sensitivity and, furthermore, suggests that different elements of the alpha range represent distinct processes (e.g. Klimesch et al., 1998).

In summary, our results clearly demonstrate that pain-free PAF is a reliable predictor of prolonged pain sensitivity. In addition to demonstrating that pain-free PAF is related to multiple models of prolonged pain, we provide compelling evidence that this relationship is stable over both immediate, i.e. minutes/hours, and more extended, i.e. weeks/months, periods of time. Furthermore, we demonstrate that pain-free PAF can be used to accurately identify high pain sensitive individuals in multiple datasets. These findings now firmly position pain-free PAF as a biomarker of pain sensitivity with untapped potential in clinical settings.

## Acknowledgements

This work was funded by an investigator-initiated research grant from Purdue Pharma L.P. to DAS. Patent pending (DAS, AM, AJF). The authors declare no other conflicts of interest.

## Supplemental Data

**Figure S1.**
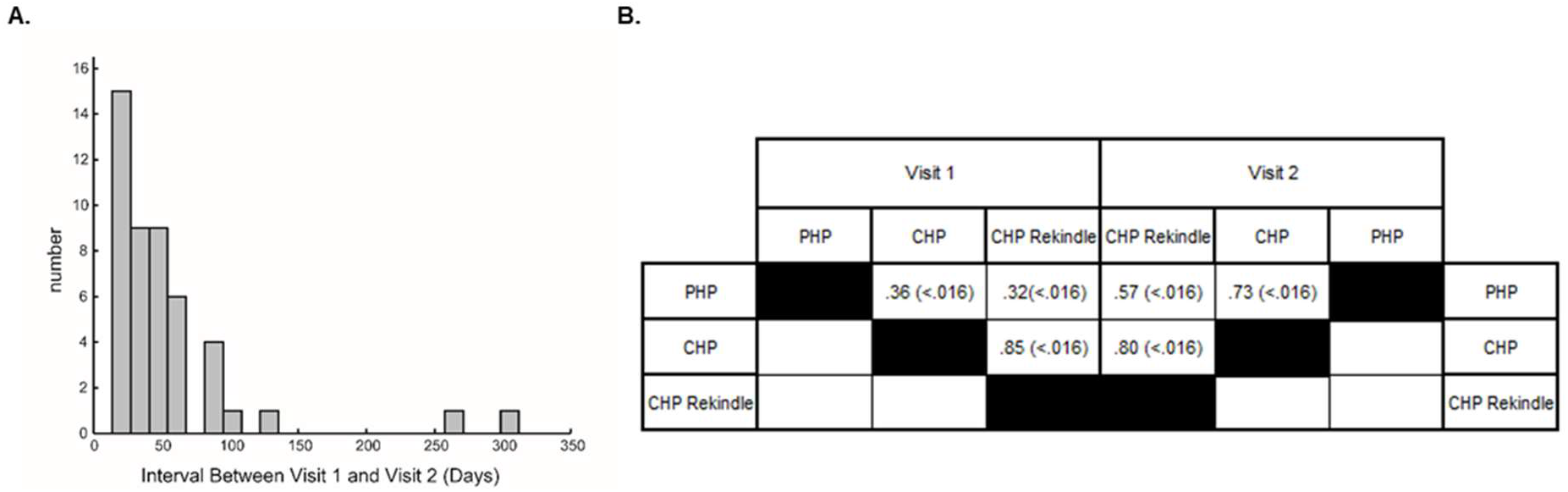
A. Histogram of days separating Visit 1 and Visit 2. On average visits were separated by 54.7 days (7.8 weeks). B. Correlations between prolonged pain tests when all participants, regardless of pain response classification, are included.

### PHP and CHP Produce Sensitization

We first sought to determine whether sensitization, a putative hallmark of prolonged pain, is present in our two prolonged pain paradigms. Inspection of the PHP time course suggest that following a decrease in pain ratings from the first to the second PHP trial, which may reflect the enhanced salience of the first stimulus train (Iannetti et al., 2008), ratings increased linearly from the second to fifth PHP stimulus train (Figure 1B). To formally test this observation, we calculated each participant’s average pain rating during PHP stimulus trains 2 and 5. Scores for all participants, regardless of pain response classification, were submitted to a linear mixed model with participants as random effects (slope included) and Visit (V1 vs. V2), Trial (2 vs. 5) and the Visit X Trial interaction as fixed effects. If PHP scores sensitize over time, then a significant main effect of Trial should be present. This analysis revealed a significant main effect of Trial, *F*_(1,88.14)_ = 19.06, *p* < .01, without a significant main effect of Visit, *F*_(1,103.08)_ = .35, *p* = .55, or significant Visit X Trial interaction, *F*_(1,88.18)_ = .02, *p* = .89. The estimated effect of Trial on PHP scores 8.61(95% Confidence Intervals: 2.13 – 15.10). This increase in scores from trial 2 (mean = 20.81, S.D. = 17.88) to trial 5 (mean = 29.31, S.D. = 21.43) in response to the same noxious stimulus is evidence of sensitization.

Two findings support the presence of sensitization during CHP. First, across all participants a pair of one-sample t-tests revealed that CHP scores were significantly greater than 0 at both V1, *t*_(57)_ = 6.63, *p* < .01, and V2, *t*_(42)_ = 5.72, *p* < .01. Second, another pair of one-sample t-tests indicated that HPTs were significantly greater than the CHP temperature, 40°C, at both V1, *t*_(57)_ = 7.90, *p* < .01, and V2, *t*_(39)_ = 8.89, *p* < .01. Pain in response to a temperature below an individual’s WDT is a strong indicator of sensitization given that we have previously demonstrated that presentation of a similar temperature without capsaicin does not produce pain (Furman et al., 2018).

**Figure S2.**
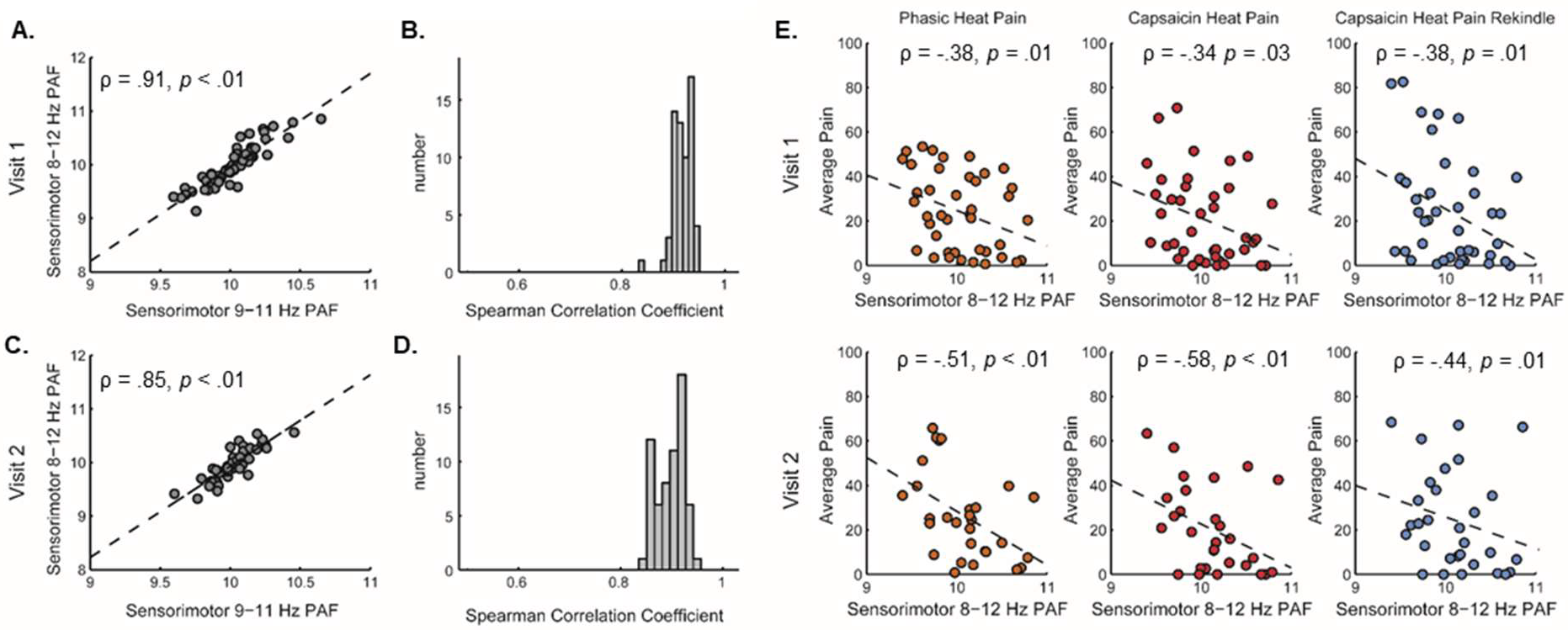
Calculating PAF using the wider 8-12 Hz range does not alter the main study conclusions. A & C. Estimates of 8-12 Hz, sensorimotor PAF and 9-11 Hz, sensorimotor PAF at both Visit 1 (A) and Visit 2 (C) are highly similar. Data comes from all participants, regardless of pain response classification, and dotted lines reflect the linear regression line of best fit. B & D. Estimates of PAF calculated using the 9-11 or 8-12 Hz range are similarly correlated at all EEG sensors. Plots reflect the distribution of Spearman correlations for 9-11 and 8-12 Hz PAF estimates across all 63 EEG channels at Visit 1 (B) and Visit 2 (D). E. 8-12 Hz, sensorimotor PAF is correlated to sensitivity to each prolonged pain test at each visit. Upper panels reflect Visit 1 pain scores and pain-free PAF; lower panels reflect Visit 2 pain scores and pain-free PAF. Presented data comes from CHP responders and High Tolerance individuals only. Dotted lines reflect the linear regression line of best fit and outliers presented in Figures 4 & 5 are omitted.

### The Frequency Range Used to Calculate PAF Does not Alter the PAF-Pain Sensitivity Relationship

Estimates of pain-free, sensorimotor PAF calculated with either a 9-11 or 8-12 Hz frequency range were highly correlated with one another whether we included all participants (V1: ρ = .91; V2: ρ = .85; Supplemental Figure S2A & S2C) or only CHP responders and high tolerance individuals (V1: ρ = .92; V2: ρ = .84). Similar correlation magnitudes between PAF estimates were evident at all scalp channels (Supplemental Figure S2B & S2D).

Correlations between pain-free, sensorimotor PAF and sensitivity to each prolonged pain test were not dramatically altered when estimating PAF with the wider 8-12 Hz range (Supplemental Figure S2E).

### Prolonged Pain Sensitivity is Similar for Men and Women

Previous studies have reported that sex may be an important variable in determining pain sensitivity (i.e. Dao & LeResche, 2000). Average pain scores (+ 1 S.D) for the sexes on each test at each visit can be seen in Supplementary Figure S3.

To determine whether sex impacted pain scores, we performed a linear mixed model for scores from each prolonged pain test with subjects as random effects and Visit (V1 vs V2), Sex (Male vs. Female), and the Visit X Sex interaction as fixed effects. Given that our study was not powered with respect to sex effects, analyses were performed on all participants regardless of pain response classification in order to maximize available statistical power. For PHP scores, this analysis revealed no significant effects of Visit, *F*_(1,41.34)_ = .37, *p* = .54, Sex, *F*_(1,53.35)_ = 1.72, *p* = .20, or Visit X Sex interaction, *F*_(1,41.34)_ = .76, *p* = .39. For CHP scores, this analysis revealed a significant Visit by Sex interaction, *F*_(1,41.86)_ = 8.95, *p* < .01, but no significant main effects of Visit, *F*_(1,41.86)_ = .06, *p* = .80, or Sex, *F*_(1,54.53)_ = 1.38, *p* = .25. For CHP rekindle scores, this analysis revealed no significant effects of Visit, *F*_(1,41.88)_ = .11, *p* = .74, Sex, *F*_(1,53.87)_ = 3.00, *p* = .09, or Visit X Sex interaction, *F*_(1,41.88)_ = 1.97, *p* > .17.

**Figure S3.**
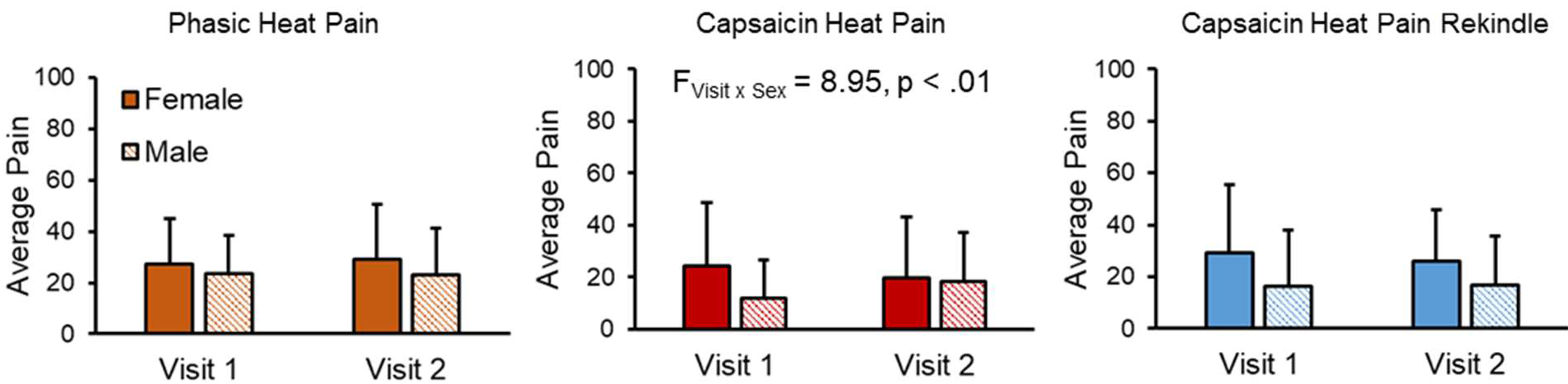
Prolonged pain scores broken by sex and visit. The only significant effect is a Visit X Sex interaction for CHP scores – scores increase for men from Visit 1 to Visit 2 and decrease for women from Visit 1 to Visit 2. Data reflect means and error bars reflect +1 standard deviation.

For CHP, the Visit X Sex interaction reflects the fact that males experience increases in CHP scores from V1 (mean = 12.10, S.D. = 14.50) to V2 (mean = 16.12, S.D. = 18.44), whereas females experience decreases in CHP scores from V1 (mean = 24.37, S.D. = 24.45) to V2 (mean = 20.01, S.D. = 23.06).

### The PAF-Pain Sensitivity Relationship is Similar for Both Sexes

One important consideration for any pain biomarker is whether it applies equally to both sexes. Correlation magnitudes for pain-free, sensorimotor PAF and CHP or PHP sensitivity were similar for both sexes; for CHP rekindle, this relationship was larger for males than females at both visits. (Supplementary Figure 4). We do not provide *p* values for these tests as our study was not powered to investigate sex differences directly.

To more formally test whether sex influences the relationship of PAF to pain sensitivity, we performed six separate moderation analyses (one for PHP, CHP, and CHP Rekindle at each visit) using PROCESS (V3.2; Hayes, 2012) implemented in SPSS. For these regression analyses, sensory test scores served as the dependent variable with pain-free, sensorimotor PAF as the independent variable and sex as a dichotomous moderator variable. As with other correlational analyses, we excluded PAF or sensory test scores greater than 2.5 SD above or below the mean value obtained at Visit 1. To account for possible multi-collinearity, independent variables and moderators were mean centered. In our moderation analyses, a significant interaction of sex and PAF would indicate that the relationship between PAF and pain sensitivity is different for the two sexes. The PAF x Sex interaction failed to reach significance for PHP scores at either V1, *t* = −.16, *p* = .87, or V2, *t* = .74, *p* = .47 or for CHP scores at either V1, *t* = −.01, *p* = .99, or V2, *t* < .01, *p* > .99. For CHP rekindle, the PAF x Sex interaction was not significant at either V1, *t* = −1.01, *p* = .32, or V2, *t* = −.98, *p* = .34. According to our moderation analyses we can conclude that the PAF-pain sensitivity relationship is not different for the two sexes.

**Figure S4.**
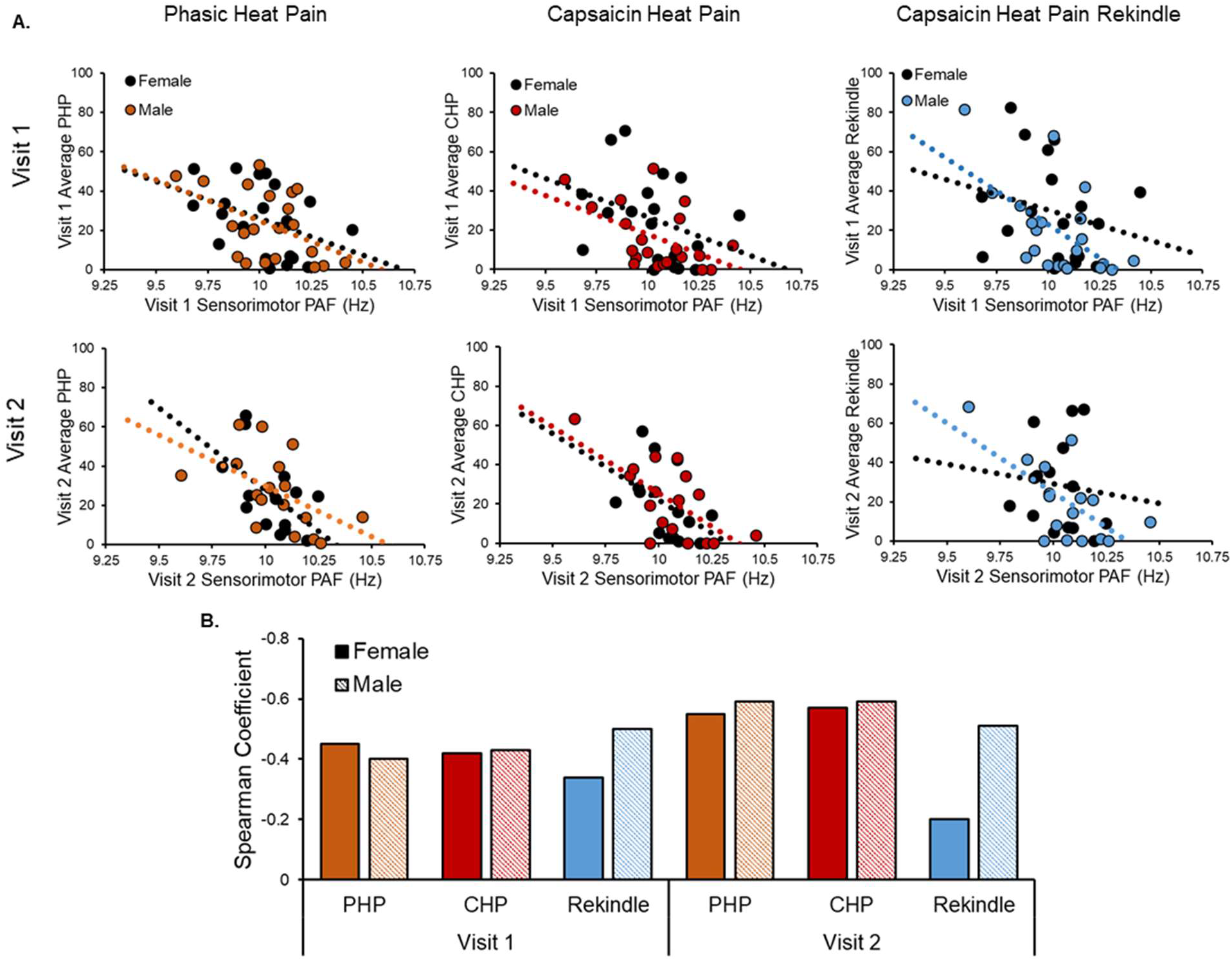
PAF bears a similar relationship to pain sensitivity for both sexes. A. Correlations between pain-free, sensorimotor PAF and prolonged pain tests are similar for both sexes at both Visit 1 and Visit 2. Note that statistical outliers presented in Figures 4 and 5 are omitted for the purpose of visual clarity. Dotted lines reflect the linear regression lines of best fit. B. Spearman correlation coefficients for the PAF-pain sensitivity relationship broken down by visit, prolonged pain test, and sex.

### The PAF-Pain Sensitivity Relationship is Evident Across the Entire Scalp

Previously unpublished findings from our lab have suggested that the PAF-pain sensitivity relationship is not privileged to channels that putatively sample the sensorimotor cortex. To determine whether similar conclusions can be drawn from the current dataset, we first calculated pain-free PAF separately at each of the 63 EEG channels. Next, we correlated PAF estimates from each channel with scores on each sensory test to yield a total of 63 correlation values for each prolonged pain test. As can be seen in Supplementary Figure S5, the distribution of sensor correlations largely recapitulated what we found when we focused only on our sensorimotor ROI. Specifically, we found that correlations between PAF and PHP, CHP and CHP rekindle were moderately large and in the negative direction. Accounting for the possible effects of volume conduction with a surface Laplacian transformation did not change these conclusions (Supplemental Figure S6).

**Figure S5.**
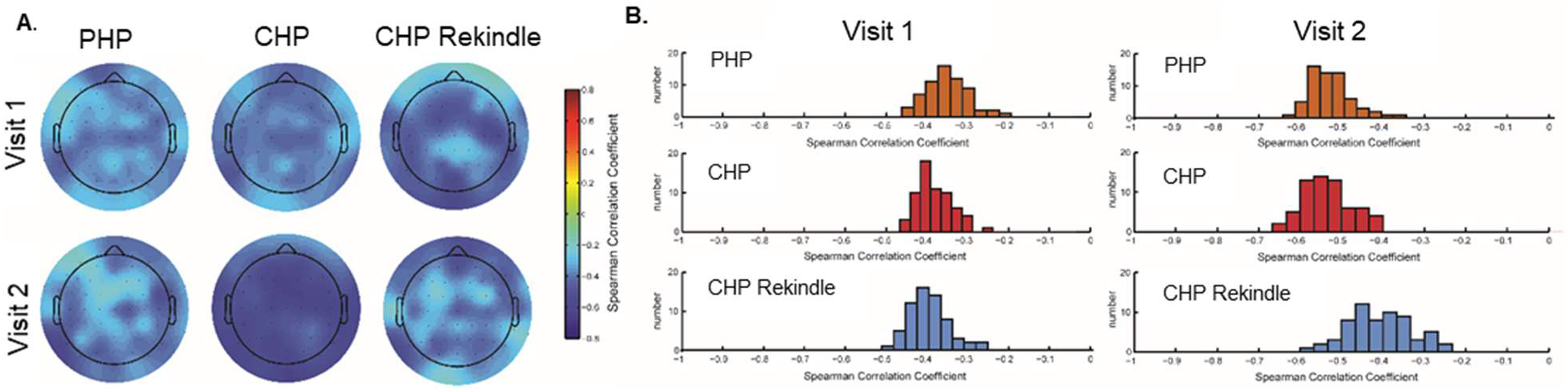
Relationship between pain-free PAF and prolonged pain sensitivity is observable across the entire EEG montage. For each prolonged pain test, and each of the 63 individual EEG sensors, the Spearman correlation between channel-level PAF and pain scores was computed to yield a total of 63 correlations coefficients for each test. Results for Visits 1 and 2 are presented either on the scalp (A) or as distributions (B).

**Figure S6.**
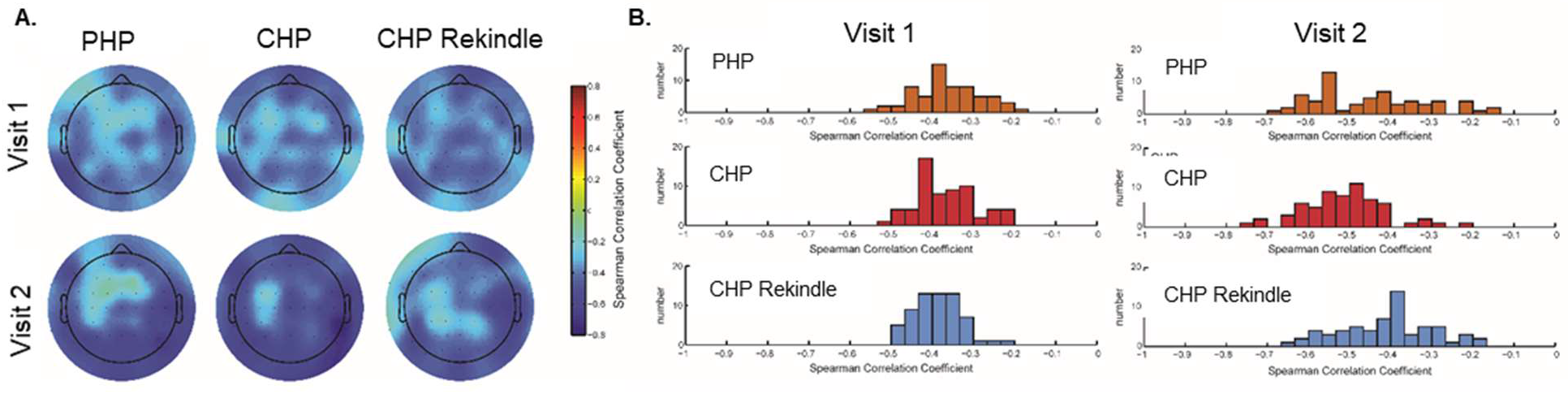
Relationship between pain-free PAF and prolonged pain sensitivity is observable across the entire EEG montage even after accounting for possible effects of voltage conduction. Prior to estimation of spectral power, a surface Laplacian transformation was applied to the preprocessed EEG data. For each prolonged pain test, and each of the 63 individual EEG channels, the Spearman correlation between channel PAF and pain scores was calculated to yield a total of 63 correlations coefficients for each test. Results for Visits 1 and 2 are presented either on the scalp (A) or as distributions (B).

### Classification of Pain Sensitivity Using PAF: Results When Including All Participants

Correlations between average pain sensitivity to all tests and pain-free, sensorimotor PAF are presented in Figure S7. Whether considering only CHP responders and high tolerance individuals (left panels), or all participants (right panels), pain-free, sensorimotor PAF was significantly related to this composite measure of pain sensitivity.

A series of within-study linear support vector machines trained on all available data from the current study identified the least sensitive individuals at above chance levels for all labelling intervals. When trying to identify the most sensitive individuals, the support vector machine was only able to do so at the smallest (10%) and largest (50%; i.e. median-split) labelling intervals. This latter result likely reflects that our composite score is an inaccurate description of the mixed sensitivity of CHP non-responders (i.e. insensitive to CHP but sensitive to PHP).

**Figure S7.**
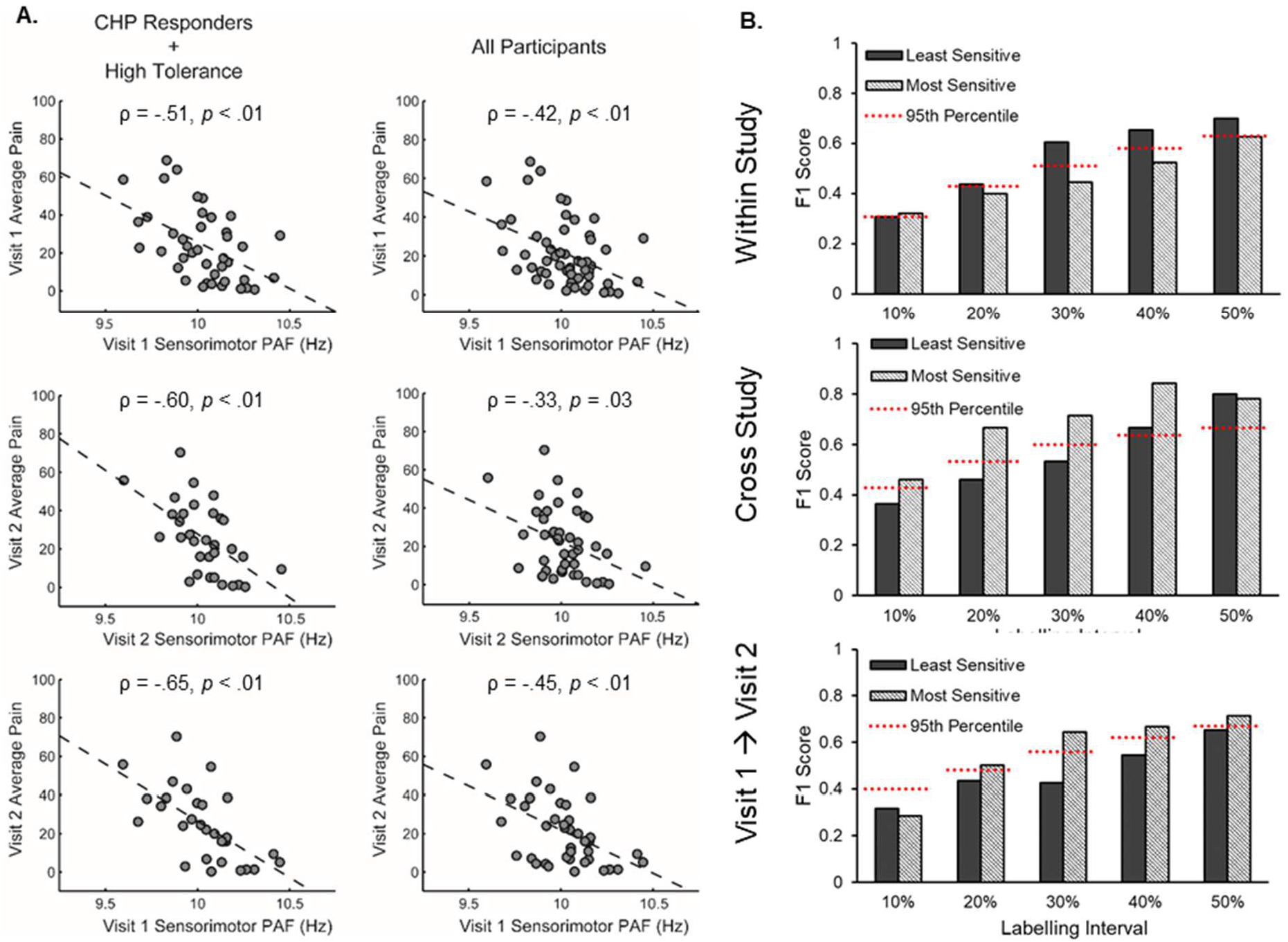
A. Pain sensitivity averaged across all prolonged pain tests is related to pain-free, Sensorimotor PAF when including only CHP Responders and High Tolerance individuals (left panels) or when including all participants regardless of pain response classification (right panels) B. Same as Figures 7B and 7D, except analyses performed while including all participants regardless of pain response classification. In brief, a support vector machine trained on Visit 1 pain-free, sensorimotor PAF predicts the identity of low pain sensitive individuals from the same study at almost all labelling intervals (top panel). A support vector machine trained on Visit 1 pain-free, sensorimotor PAF predicts the identity of high pain sensitive individuals from an independent study at all labelling intervals (Furman et al., 2018; middle panel). A support vector machine trained on Visit 1 pain-free, sensorimotor PAF predicts the identity of Visit 2 high pain sensitive individuals at almost all labelling intervals (bottom panel). An F_1_ score of 1 indicates perfect classifier performance and the dashed red lines reflect the 95^th^ % of a null distribution of F_1_ scores.

A cross-study linear support vector machine trained on all available data was able to identify the most sensitive individuals from a separate dataset (Furman et al, 2018) at all labelling intervals. For the least sensitive individuals, this support vector machine only performed at above chance levels for the two largest labelling intervals.

